# An Oscillatory Shear Index-based Model to Describe Progressive Carotid Artery Stenosis

**DOI:** 10.1101/2022.02.04.479106

**Authors:** Peter Patalano

## Abstract

Background and aims: This study describes and demonstrates the applicability of a novel *in silico* method for modeling progressive carotid artery stenosis using the oscillatory shear index (OSI) as the basis of stenosis. Methods: Three-dimensional reconstructions of 11 carotid arteries were generated using patient-derived magnetic resonance angiography and duplex ultrasound data. Computational fluid dynamic simulations were sequentially generated following computational stenosis assessment, and corresponding changes in OSI were observed and used as measure of morphological stabilization. Results: Six carotid models showed progressive stenosis with statistically significant increases in regions of high OSI (OSI > 0.2, p < 0.05) with eventual carotid occlusion in one of the cases. Three models remained free or nearly free of increased OSI, whereas one model showed an overall decrease in high OSI regions (p < 0.05) and another trended in that direction but did not achieve statistical significance (p = 0.145). Conclusions: To our knowledge, this is the first computational model describing progressive stenosis in any peripheral artery including the carotid. Taken together, this study provides a novel framework for computational hemodynamic investigations on progressive atherosclerosis in the carotid artery.

## Introduction

Atherosclerotic plaques near the carotid bifurcation, particularly in the internal carotid artery (ICA), are believed to be the most common detectable cause of ischemic stroke. When classified as high grade, asymptomatic patients have 4.6–18.5% risk of experiencing stroke within 5 years [1]. Carotid endarterectomy and stenting are common treatment options for symptomatic and severe asymptomatic carotid artery stenosis [2]. Although progression of stenosis (regardless of treatment) cannot be reliably predicted, it remains a clinically relevant problem. Hemodynamic factors related to low wall shear stresses (WSSs) have been implicated in the development of atherosclerosis within the carotid artery [3, 4, 5]. However, it is still difficult to predict which patients will progress and which will stabilize, as the available approaches are based on catch-all efforts to monitor patients with moderate stenotic lesions using repeated imaging data.

WSSs, which describe the forces acting in parallel to the vessel wall, is a well-studied factor involved in the genesis of atherosclerotic plaques in the arterial system. Areas of high WSS tend to result from highly organized laminar flow, whereas areas of low WSS tend to be characterized by complex flow patterns (including oscillatory flows). Low WSS has been implicated in the activation and deactivation of inflammatory pathway genes that, when left unchecked, result in the triggering of the atherosclerotic cascade in both in vitro and in vivo studies [6, 7, 8, 9].

Framing questions concerning aberrant fluid patterns has paved the way for investigations aiming to test numerical and computational methods for a better understanding of this problem. Using computational fluid dynamic (CFD) approaches and the finite element method (FEM), it became possible to model complex fluid flows in the arterial lumen by reconstructing three-dimensional (3D) models from imaging data, such as computer tomographic angiography (CTA) and magnetic resonance angiography (MRA). Additionally velocity data can also be obtained from duplex ultrasound (DUS) and certain MR sequences, which can also aid the modeling and simulation process [10, 11].

Although WSS metrics were reported to be strongly associated with the initiation and propagation of atherosclerotic plaques, spatial prediction of plaque development remains difficult, owing partially to challenges in reproducibility [12, 13]. Prior studies have focused on correlations between WSS and the spatial location of the plaques; however, the hemody namic effects of modeled stenosis may have on future CFD simulations remains unclear. Hence, this approach provides only a static picture of the development and progression of atherosclerosis.

The present study explores the hemodynamic changes that are induced by modeled stenosis to generate a model for atherosclerosis progression. A dynamic modeling approach was used for a better understanding of the effects of *in silico* stenosis on future hemodynamics and subsequent progression of stenosis. The problem was addressed in an iterative manner, where flow in a native artery was simulated and stenosis was generated according to the corresponding OSI. This process was repeated, while considering any hemodynamic changes induced by *in silico* stenosis generated by previous simulations. Thus, rather than attempting to predict point-based atherosclerosis, the proposed new method focuses on the effect of *in silico* stenosis on future hemodynamics and subsequent changes in OSI. A central question is raised of whether stenosis in a region of high OSI leads to more favorable or less favorable hemodynamics in future CFD simulations. The luminal surface was treated as a changing landscape, as new stenoses arise, the flow of fluid through the lumen is necessarily changed. The question can then be rephrased - does *in silico* stenosis serve to stabilize or destabilize future flows?

Blood flow in the human carotid artery is governed by the physical properties of the fluid (i.e. viscosity, density, among other characteristics), the applied force that acts on it (i.e.heart), and the boundaries with which the blood interacts(i.e. vessel walls). As with all other fluids, blood flow is governed by the principles of mass, momentum, and energy conservation, which are in turn physical laws that can be represented by the incompressible Navier-Stokes equations,which describe viscous fluid flow in 3D. (Eq. 1).

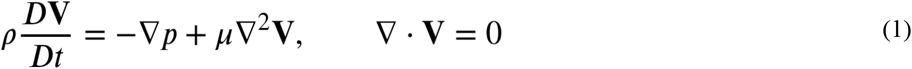

Equation 1 states that the acceleration 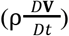 in a fluid is equal to the sum of the pressure forces (−**∇***p*) and the viscous diffusion forces (*μ***∇**^2^**V**). Additionally, in the incompressible form, the continuity relationship (**∇**·**V**= 0) shows that the amount of fluid that flows into a volume must be equal to the amount that flows out. Solving this system of partial differential equations allows us to understand the velocity and trajectory of blood within an artery, as well as the forces that are imparted by the fluid on the vessel walls for a fluid of known density (ρ) and viscosity (*μ*). These Principles are often applied experimentally to determine the stability of implanted devices, as well as to better understand the effects of complex flows on the vascular biology and gene expression related to pathological processes, such as atherosclerosis and aneurysm formation [14, 15, 16, 17, 18].Experimental research in medicine and bioengineering have led to the discovery of certain complex flows that serve as the initiators and propagators of both aneurysms and atherosclerotic disease [19, 7, 8, 20]. Of specific interest to our work are flows that result in layer separation and oscillatory patterns at the carotid artery bifurcation. Since these flow patterns and related WSS have been implicated in atherosclerosis, being able to model and quantify flows in this region may have important implications in the prediction of atheroscleroticplaque formation and progression of clinically significant stenosis.

Low and oscillatory wall stresses in animal models have been shown to promote gene expression which facilitates the cascade of vessel wall inflammation, endothelial proliferation, apoptosis, permeability, and cholesterol deposition. Similar observations have been made in clinical studies. Indeed, oscillatory flow patterns in particular have been implicated in the progression of atherosclerosis [21, 12].

Various indices have been numerically developed and experimentally verified to quantify these stresses and their associated flow patterns. The one we found most useful was the OSI, owing its association with plaque progression in clinical studies [22, 23]. At a given point on the luminal surface, for a given time period, OSI (Eq. 2) quantifies how much wall shear stress (τ_*w*_) deviates from the average value at that point. Wall shear stress (τ_*w*_) is the force imparted by a fluid of viscosity, *μ* moving parallel to the wall with velocity, *u* at distance, *y* from the wall. OSI describes whether flow at a certain point tends to be laminar and simple (OSI = 0) or if it is subjected to maximal amounts of flow reversal (OSI = 0.5) [24].

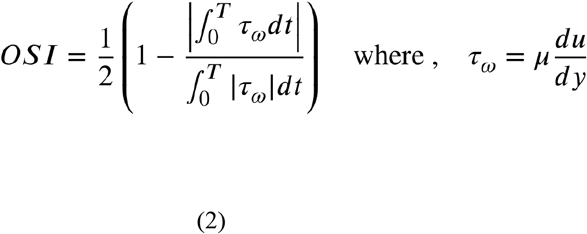

This index was used as the basis of a new *in silico* stenosis model.

After running the fluid dynamic simulation for sufficient time to achieve flow stabilization, OSI was calculated across the luminal surface. Fluid flow was first simulated then each element (*P*_*i*_) on the vessel lumen was translated by the magnitude of the corresponding *OSI*_*i*_ in the direction of the inward facing the normal surface (*P*_*i*_)(Figure 2), resulting in a novel luminal surface and *in silico* stenosis. Fluid dynamic simulation was then repeated on the novel surface and observed visually for new areas of high OSI. It was our hypothesis that at least two situations could emerge after each CFD simulation and the *in silico* stenosis generated:

**Figure 1:**
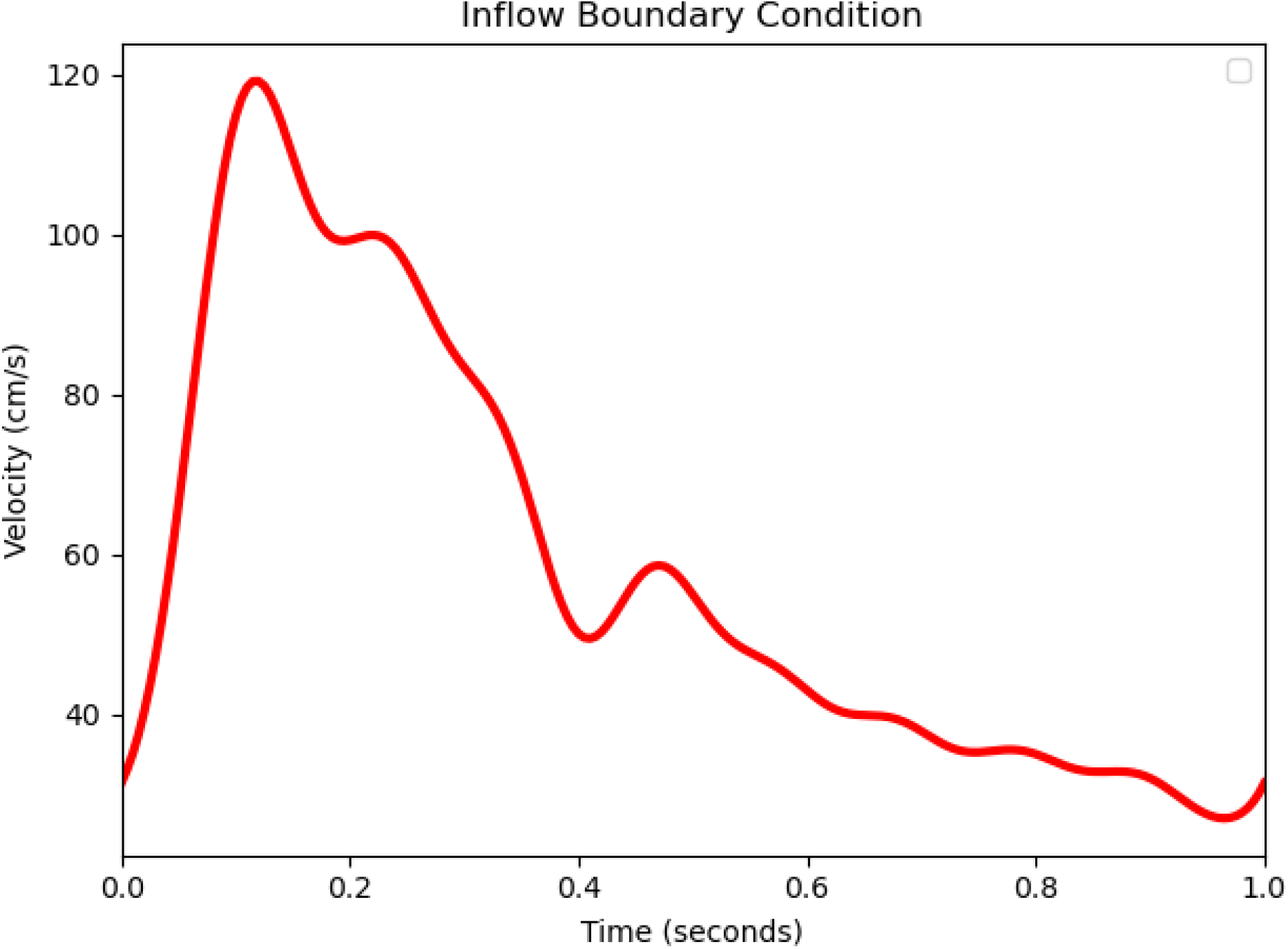
Example of the inflow boundary condition used for computational fluid dynamic simulations. Peak systolic velocity(PSV) is obtained from duplex ultrasound (DUS) data and the waveform is estimated using the feature points seen in the human common carotid waveform [25].

**Figure 2:**
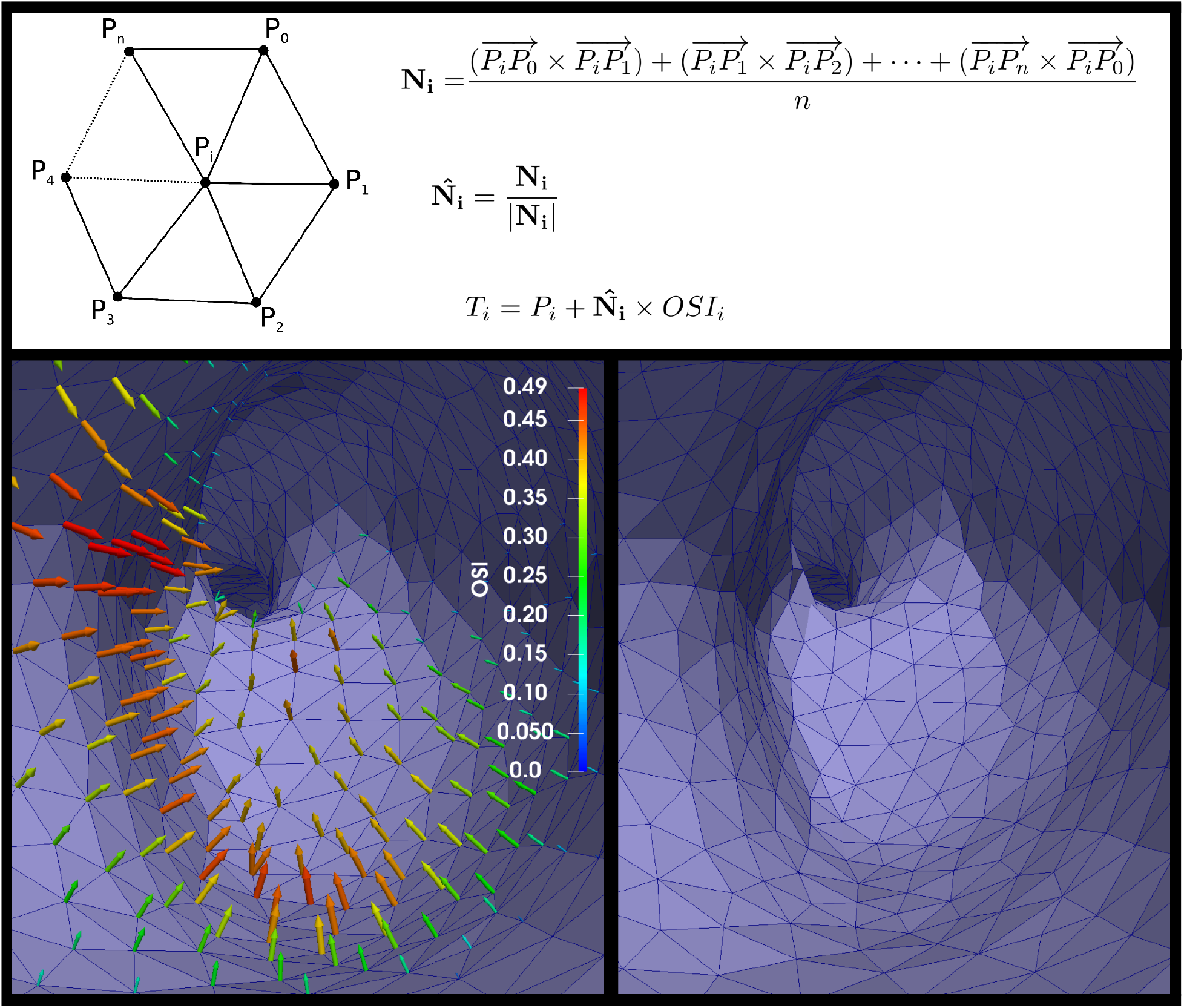
The top panel describes the stenosis generation process. Following each simulation, after OSI is calculated, the inward facing normal, N_i_ is calculated for a given point (*P*_*i*_) by averaging the normal direction of the adjacent faces. The unit normal, N(hat)_i_ is then calculated. The translation, *T*_*i*_ is then generated by moving the point *Pi* in the direction of the normal by the magnitude of OSI. The bottom panels graphically demonstrate the process of stenosis generation. The panels show the luminal surface of an arbitrary vessel. The arrows face inwawrd at each point, their length and color correspond to the magnitude of OSI at that point. The bottom right panel shows the result of moving each point to the tip of each arrow, thus completing the translation and stenosis.

1. the stenosis function could lead to stabilization of OSI,resulting in less stenosis on subsequent simulations, or
2. the stenosis function could lead to new areas of highOSI, propagating the stenosis on subsequent simulations.

Should situation (1) occur, subsequent analyses would yield no meaningful stenosis, whereas in situation (2), the stenosis would propagate until situation (1) was encountered or the luminal surface arrives in apposition, i.e., an occlusion occurs. Thus, this iterative process can be used to model patient-specific progression of stenosis, a technique which may have both experimental and clinical applications in the future.

## Methods

This study was conducted under the guidance of the NYU School of Medicine Institutional Review Board. MRA and DUS data were obtained from the medical record of patients being followed for suspected carotid artery stenosis. Patients were included if an MRA and DUS of the carotid arteries were obtained at the time of initial consultation. None of the MRA exams contained velocity data and thus DUS velocities were used for this purpose. All Studies were made anonymous after acquisition to protect patient information.

Patient imaging data was first anonymized and assigned a random 3 digit ID number. DICOM images were loaded into the Vascular Modeling Toolkit (VMTK, http://www.vmtk.org) and image segmentation was achieved using built-in semi-automatic methods (colliding fronts and fast marching algorithms). 3D surfaces were obtained using implementation of the marching cubes algorithm of the VMTK. The surfaces were then clipped and flow extensions were added to the common carotid artery (CCA), ICA, and external carotid artery (ECA). The bounding boxes used to clip the branches were stored as multiblock surfaces. 3D surfaces were converted into polygonal meshes using SimVascular(SV, http://www.simvascular.org) with increased density of elements at the boundary wall.

Due to the lack of detailed velocity data over the cardiac cycle in the CCA, a pulsatile waveform estimation was generated to simulate pulsatile inflow. PSV at the CCA was first obtained from DUS.Additional points were then generated based on the relative amplitude of feature points seen in the CCA waveform in older adults. The estimated EDV obtained by this method was then compared with DUS data. We arbitrarily chose a difference of 10cm/s between estimated and observed EDV to be acceptable (Table 1). Using the built-in Fourier estimation of SV, the inlet velocity based pulsatile waveform was then generated.

**Table 1:**
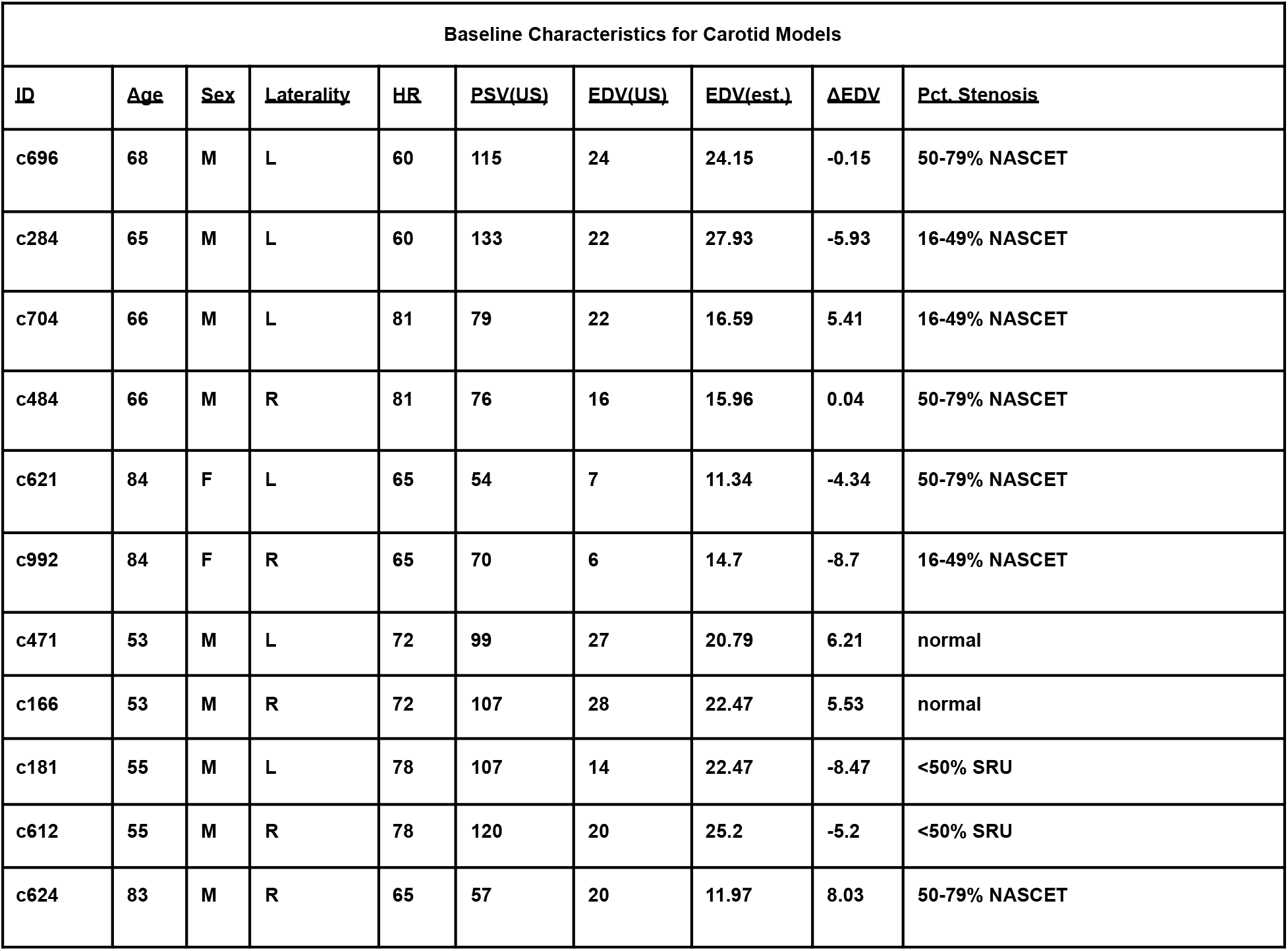
Baseline characteristics from patient-derived data including the reported stenosis percent as recorded with duplex ultrasonography.EDV (est.) is the EDV which was used in the pulsatile waveform for simulation approximated with pulse waveform estimation.

| Baseline Characteristics for Carotid Models |  |  |  |  |  |  |  |  |  |
| --- | --- | --- | --- | --- | --- | --- | --- | --- | --- |
| ID | Age | Sex | Laterality | HR | PSV(US) | EDV(US) | EDV(est.) | ΔEDV | Pct. Stenosis |
| c696 | 68 | M | L | 60 | 115 | 24 | 24.15 | -0.15 | 50-79% NASCET |
| c284 | 65 | M | L | 60 | 133 | 22 | 27.93 | -5.93 | 16-49% NASCET |
| c704 | 66 | M | L | 81 | 79 | 22 | 16.59 | 5.41 | 16-49% NASCET |
| c484 | 66 | M | R | 81 | 76 | 16 | 15.96 | 0.04 | 50-79% NASCET |
| c621 | 84 | F | L | 65 | 54 | 7 | 11.34 | -4.34 | 50-79% NASCET |
| c992 | 84 | F | R | 65 | 70 | 6 | 14.7 | -8.7 | 16-49% NASCET |
| c471 | 53 | M | L | 72 | 99 | 27 | 20.79 | 6.21 | normal |
| c166 | 53 | M | R | 72 | 107 | 28 | 22.47 | 5.53 | normal |
| c181 | 55 | M | L | 78 | 107 | 14 | 22.47 | -8.47 | <50% SRU |
| c612 | 55 | M | R | 78 | 120 | 20 | 25.2 | -5.2 | <50% SRU |
| c624 | 83 | M | R | 65 | 57 | 20 | 11.97 | 8.03 | 50-79% NASCET |

In individual models, pulsatile blood flow was simulated.The arterial wall was assumed to be rigid. Blood was assumed to be an incompressible, Newtonian fluid with density ρ=1.06*g*·*cm*−3 and viscosity *μ*=0.004*Pa*·*s*. The fluid dynamic solver from SV was used for fluid flow simulations and was solved using the High Performance Computing Cluster at NYU. At the inlet boundary, we simulated velocity-based pulsatile flow with a parabolic velocity profile. Initial Velocity during the cardiac cycle was estimated based on the waveform feature points, as described above, and based on patient-specific clinical DUS PSV at the CCA. Resistance boundary conditions were used at the outlets. Other Boundary conditions included rigid-walls and zero-slip. The simulation was run for a total of six cardiac cycles based on the baseline heart rate of the patient. A total of 200 time points per each cardiac cycle were simulated.

Following CFD simulation, the *in silico* stenosis was generated using ParaView (PV, http://www.paraview.org) and our own code written in Python using the VMTK and VTKpackages. First, flow extensions were removed using the previously stored clipping box data. Next, inward facing normal units were generated at each point on the polygonal dataset and point-wise translation was completed. (Figure 2). The resulting stenosis was first visually verified, then quantified based on the stability of OSI. To assess for OSI stability following each iteration of stenosis, the variance of the OSI was calculated across the luminal surface and was compared with subsequent runs using Levene’s test. In a vessel that is undergoing flow stabilization, we expect variance to become smaller and approach zero. We therefore used Levene’s test to assess whether a change in variance was statistically significant or not. We considered a p-value of < 0.05 to be statistically significant. We chose an OSI cutoff of > 0.2 to be considered high, based on previous reports showing increased risk of atherosclerosis of OSI between 0.1 and 0.3 [26, 27, 28].

## Results

The models were first visually inspected between simulations and obvious luminal narrowing was noted, especially in those with relatively large variance of OSI. As expected,each carotid model deformed to a certain degree with each subsequent simulation.

It was noted that the distribution of OSI could change in the vessel lumen without creating regions of high OSI (OSI> 0.2), as observed in c181. The variance of OSI across the luminal surface showed a statistically significant increase from 0.000394 to 0.000592 (*p* <<0.001) without introducing regions of high OSI. A similar case arose in model c612, which showed a statistically significant increase in variance without generating regions of high OSI (*p* <<0.001).

After a single simulation, the morphological changes in 624 were drastic enough to warrant further investigations. Three more simulations were run and the analysis failed dueto complete lack of flow at the internal carotid outlet in the final simulation, representing a total occlusion of the early ICA (Figure 3).

**Figure 3:**
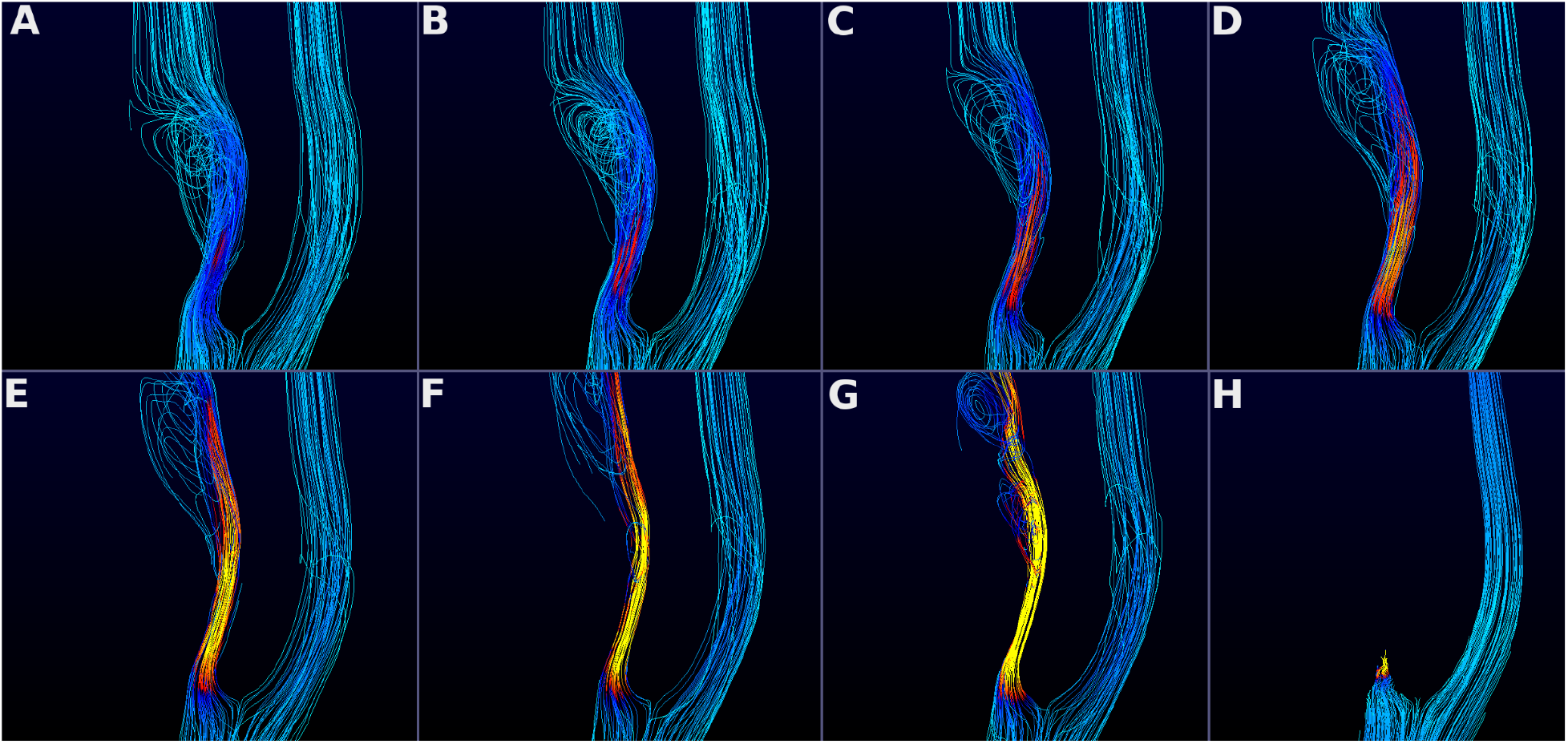
A particle trace in sequential simulations and stenoses in model c624. Cooler and warmer colors represent lower and higher relative velocities, respectively. The top left panel (A) represents the computational fluid dynamic simulation based on the native magnetic resonance imaging geometry. Early stages of the stenosis progression process is shown in (B), (C), and(D). (E), (F) and (G) further show that the stenosis continues to worsen and leads to a total occlusion (H) at the proximal internal carotid artery.

A modest increase (< 1%) in high OSI regions was observed in c992 (3.09% to 3.46%, *p*=2.42×10^−6^), c471 (2.59% to 3.21%, *p*=0.0121) and c621 (5.49% to 5.94%, *p*=0.0218) (Table 2). Larger increases in high OSI regions were also detected in c166 (0.49% to 2.58%), c284 (8.9% to 13.44%), and c624 (5.02% to 8.8%), all of which reached statistical significance. These observations further supported the hypothesis that regions with high OSI values predispose arteries to atherosclerosis and associated luminal narrowing.

**Table 2:**
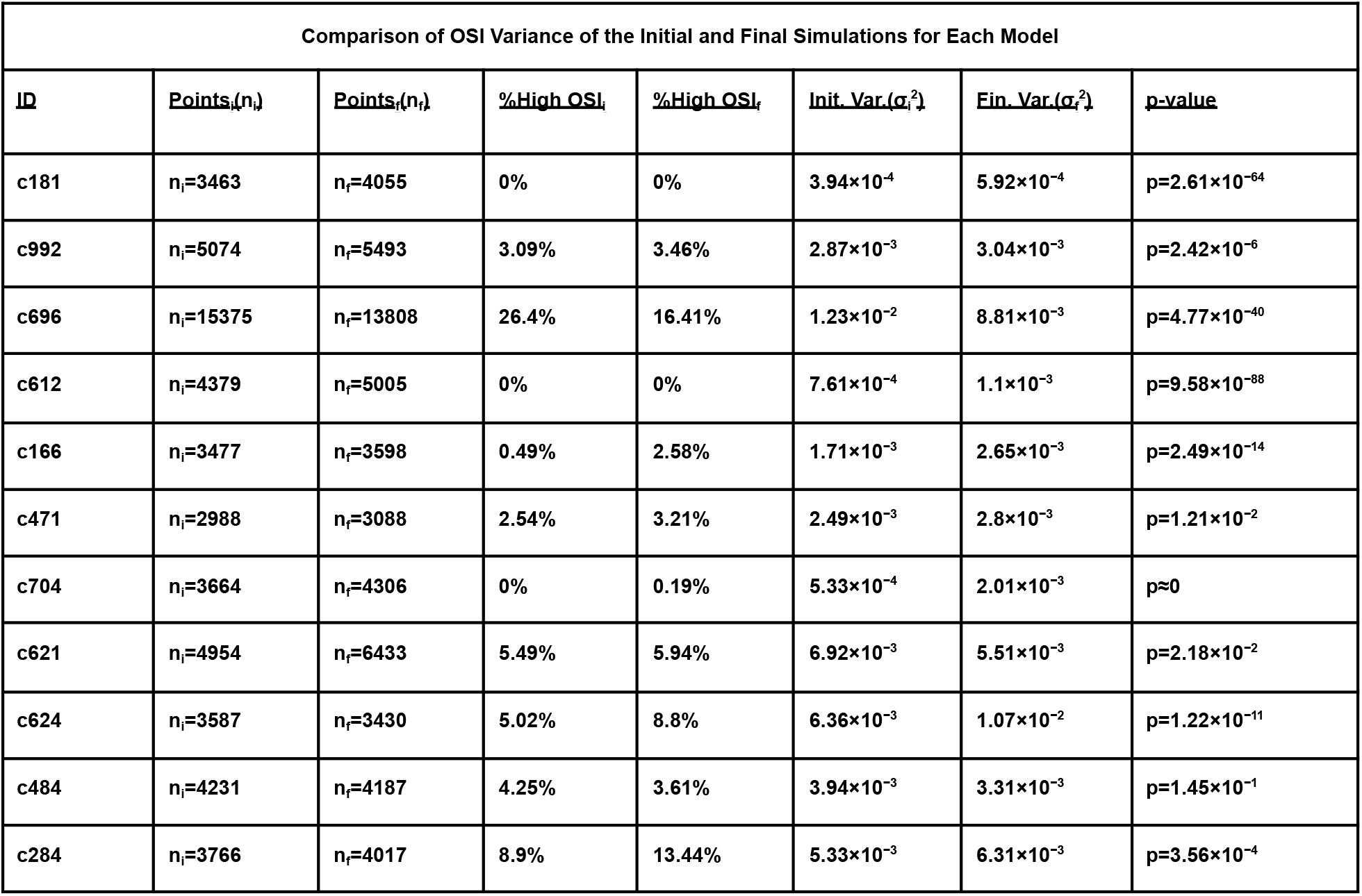
The table shows the percent increase in regions of high OSI between the initial and final simulation, as well as the associated change in variance of OSI between initial and final simulations. P-values were obtained by comparing the initial and final variance of OSI using Levene’s test.

One model (c696) underwent morphological changes that led to an overall decrease in high OSI regions (26.40%to 16.41%,*p* <<0.0001) whereas another model (c484) trended in that direction but did not achieve statistical significance (4.25% to 3.61%,*p*= 0.145) (Table 2). Qualitatively, these were seen as changes from bulbous morphologies to those that were more cylindrical.

We grouped these changes into two patterns due to the downstream morphological consequences on subsequent fluid dynamic simulations. (1) A statistically significant decrease, or no statistically significant change in OSI variance between simulations; or (2) a statistically significant increase in OSI variance between simulations. The latter category could be further refined by observing models with substantially elevated OSI (OSI > 0.2), which may suggest risk of atherosclerotic progression.

## Discussion

Although low WSS and high OSI have been implicated in the progression of atherosclerosis, their predictive utility has remained limited [13]. This may be partially due to the challenge in predicting *in vivo* stenosis from computationally acquired hemodynamic metrics as well as the fact that previous models reach spatial plaque prediction following a single simulation. Our methods may help overcome such limitations as we approach stenosis as an ongoing interplay between atherosclerosis and alterations in complex fluid flows. This research presents a dynamic landscape of aberrant fluid flows and highlights the effects of modeled stenosis on future fluid flows. This approach is the first to offer computational insight into an artery undergoing progressive stenosis, addressing whether the stenosis has a stabilizing or destabilizing effect on the flow patterns.

An example in which stenosis may stabilize flow can be seen in c696 (Figure 4). Although this model underwent stenosis, it showed an overall decrease in OSI variance. A similar trend is shown in c484, a model that was predicted to undergo further stabilization in later simulations.

**Figure 4:**
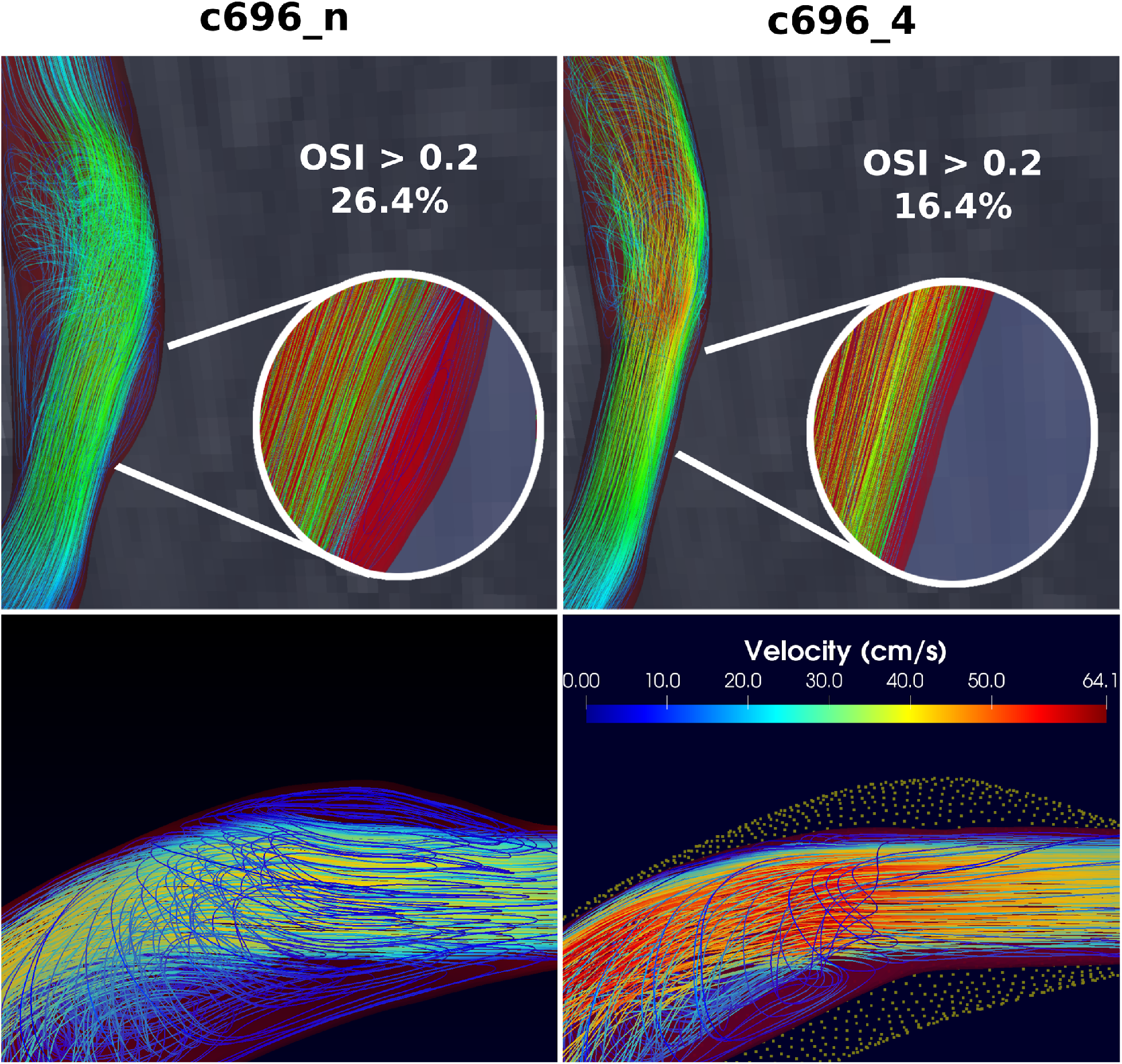
Representative example of stabilizing stenosis. The top panels show the native flow and following four iterations of stenosis with their associated percentage of luminal surface with high oscillatory shear index (OSI). Noteworthy is the resolution of the more disorganized fluid flow. The bottom panels shows the portion of the internal carotid artery (ICA) with native flow (bottom left) and following stenosis (bottom right). Additionally, the bottom right panel shows an overlay of the previously patent portion of the artery with a stabilizing flow pattern being noticeable across the region of stenosis.

Conversely, the effect of stenosis may be destabilizing, as was observed in the other models, which showed an overall increase in OSI variance and, in some instances, a drastic increase in regions of high OSI (Figure 5). Thus, it is reasonable to consider that fluid flows represent an ever-changing landscape in the vessel lumen. As areas of high OSI emerge and undergo stenosis, new areas of high OSI may develop and even disappear. This study outlines and demonstrates a novel computational method for modeling the progression of stenosis at the carotid artery bifurcation based on OSI. Additionally, we offer an approach for quantifying its success and have demonstrated encouraging preliminary results that have directed our attention toward understanding atherosclerosis as a dynamic process modulated by the interplay of plaque geometry, vessel morphology, and hemodynamics based on the effects of the resulting stenoses.

**Figure 5:**
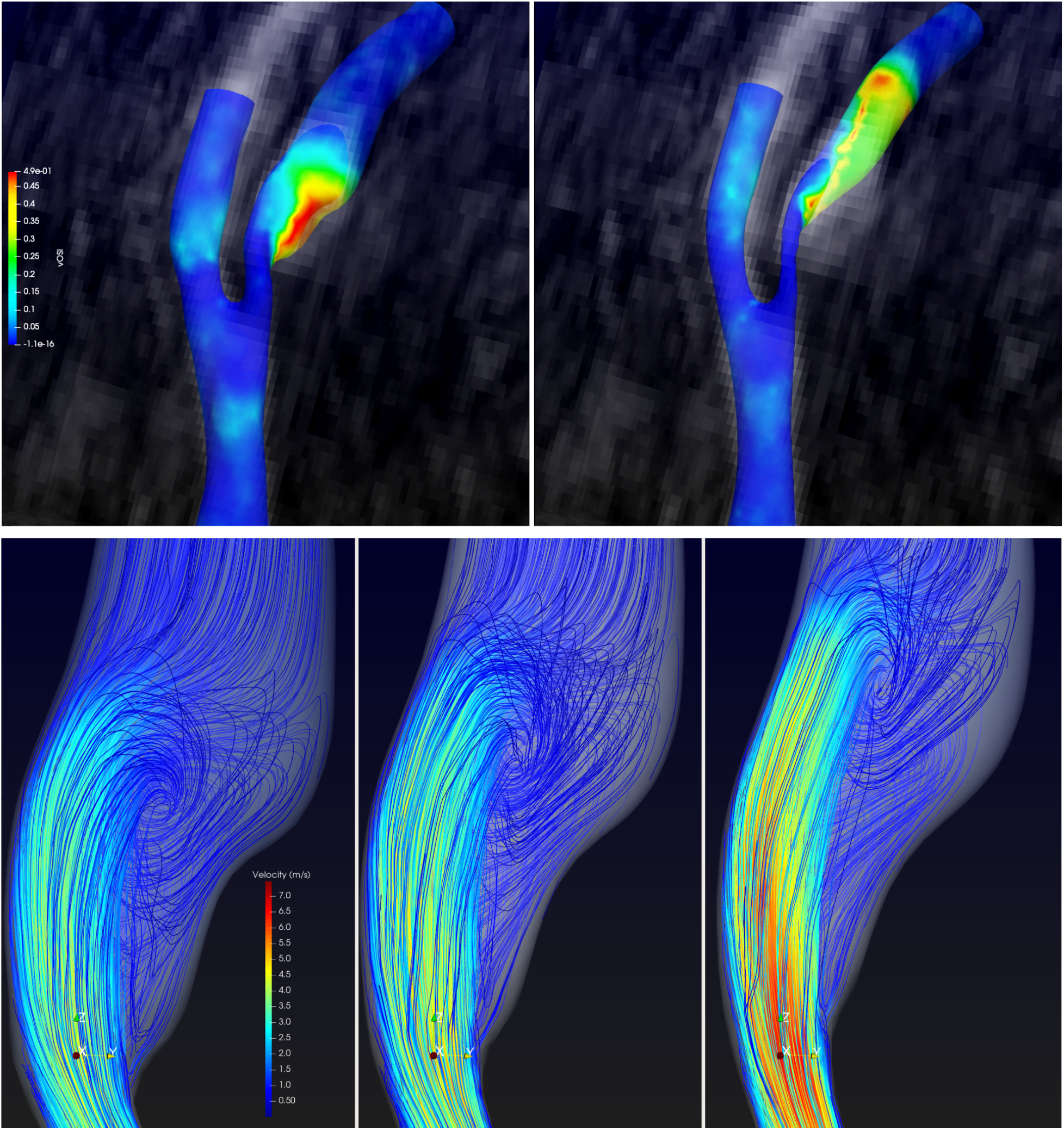
Impact of destabilizing stenosis on the surface and intraluminal space. The top panels show the luminal surface distribution of oscillatory shear index (OSI) before and after an iterative stenosis was generated. Warmer and cooler colors represent higher and lower OSI, respectively. Notice the migration of the region of high OSI more distally along the internal carotid artery (ICA). The bottom panels show the distal migration of unorganized fluid flow as the stenosis progresses. Warmer Colors represent higher velocities. As expected, the velocity across the lesion increases as the stenosis progresses.

The concept that flow disturbance is an important component in the initiation and progression of stenosis has been demonstrated several times. Specifically, arterial bifurcations throughout the vascular system are prone to the generation and progression of atherosclerosis, which eventually may lead to clinically significant stenosis. The vessel branching leads to flow instability through low and oscillatory stresses, which initiate and propagate the atheroscleroticcascade in at-risk individuals [19, 4, 6, 7]. Despite the robust body of research relating arterial bifurcations with aberrant flow leading to atherosclerosis, their role in predictive modeling remains unclear.

The methods described in this study may provide new foundations to begin modeling progressive stenosis as the methods are refined and validated against *in vivo* data. There are unresolved questions regarding the validation of the OSI > 0.2 that was used to describe high OSI regions. This value was chosen based on biologically informed reasoning and merely used as a quantitative indicator of destabilization; however, a more context specific nuance could be introduced by using a threshold value for OSI, above which selective stenosis could be generated.

One limitation of our approach stems from the stenosis function itself. Although boundary layer thickness and wall stress calculations consider the vessel radius, our stenosis function does not. This may overestimate stenosis for smaller vessels and regions containing existing tight stenosis. As the main purpose of this paper is to demonstrate the effect of *in silico* stenosis on future flows and the size similarity in the human carotid, we did not correct for this. We do not predict substantial deviations over the course of a few simulations but will focus on identifying and developing appropriate radius based normalization in future efforts.

As mentioned above, high OSI values were considered to be > 0.2 based on evidence that suggests that elevated OSI leads to atherosclerotic-related gene expression and consequent increased risk of atherosclerosis [27, 28]. Therefore,we would expect that defining an OSI threshold for generating stenosis may lead to earlier morphological stabilization. For example, in c181 and c612, 0% of the luminal surface was found to have an OSI> 0.2. If the threshold at which stenosis is generated is set to 0.2, we would expect no morphological change of the surface on subsequent simulations. Similarly, in models with regions of concentrated high OSI, we expect more focal stenoses along with possible changes in flow which could be more similar to the vascular remodeling response seen *in vivo*[29].

While questions such as these are resolved or further refined and further analyses are conducted, we hope to develop an efficient computational method centered around the following main questions: (1) Does threshold modeling play a role in the propagation of *in silico* stenosis? (2) In at-risk patients, are there certain geometric configurations that lead to inevitable progression of stenosis? (3) Can we identify specific features that make these vessels unique?

Using these questions, we hope to refine and validate this approach with the aim of creating a clinically applicable tool. An improved ability to predict the progression of stenosis in a patient-specific manner may represent a profound leap forward in both areas of clinical decision making as well as unraveling the complex relationship between biology, vessel morphology, and fluid dynamics.

## Acknowledgements

Special thanks to Caron Rockman, MD, for her support and sponsorship of this research, as well as GlennJacobowitz, MD, and the Division of Vascular Surgery at NYU. Additionally, the author would like to acknowledge the developers of SimVascular, VMTK, and ParaView forallowing the open-source usage of their code.

## References

[1] D. Inzitari, M. Eliasziw, P. Gates, B. L. Sharpe, R. K. Chan, H. E. Meldrum, and H. J. Barnett. The causes and risk of stroke in patients with asymptomatic internal carotid artery stenosis. North American symptomatic carotid endarterectomy trial collaborators.The NewEngland Journal of Medicine, 342(23):1693–1700, June 2000. ISSN0028-4793. doi: 10.1056/NEJM200006083422302.

[2] Thomas G. Brott, Robert W. Hobson, George Howard, et al. Stenting versus endarterectomy for treatment of carotid-artery stenosis.New EnglandJournal of Medicine, 363(1):11–23, July 2010. ISSN 0028-4793,1533-4406. doi: 10.1056/NEJMoa0912321. URLhttp://www.nejm.org/doi/abs/10.1056/NEJMoa0912321.

[3] Alexander M. Nixon, Murat Gunel, and Bauer E. Sumpio. The critical role of hemodynamics in the development of cerebral vascular disease: A review.Journal of Neurosurgery, 112(6):1240–1253, June 2010. ISSN 0022-3085, 1933-0693. doi: 10.3171/2009.10.JNS09759. URLhttps://thejns.org/view/journals/j-neurosurg/112/6/article-p1240.xml.

[4] Saurabh S. Dhawan, Ravi P. Avati Nanjundappa, Jonathan R. Branch, et al. Shear stress and plaque development.Expert Review of Cardiovascular Therapy,8(4):545–556, April 2010. ISSN 1744-8344. doi: 10.1586/erc.10.28.

[5] Umberto Morbiducci, Annette M. Kok, Brenda R. Kwak, Peter H. Stone, David A. Steinman, and Jolanda J. Wentzel. Atherosclerosis at arterial bifurcations: evidence for the role of haemodynamics andgeometry.Thrombosis and Haemostasis, 115(3):484–492, March2016. ISSN 2567-689X. doi: 10.1160/TH15-07-0597.

[6] Kyung-Sun Heo, Keigi Fujiwara, and Jun-ichi Abe. Shear stress and atherosclerosis. Molecules and Cells, 37(6):435–440, June 2014.ISSN 0219-1032. doi: 10.14348/molcells.2014.0078.

[7] Jeng-Jiann Chiu, Shunichi Usami, and Shu Chien. Vascular Endothelial responses to altered shear stress: Pathologic implications for atherosclerosis.Annals of Medicine, 41(1):19–28,January 2009. ISSN 0785-3890, 1365-2060. doi: 10.1080/07853890802186921. URLhttp://www.tandfonline.com/doi/full/10.1080/07853890802186921.

[8] Michael A. Gimbrone and Guillermo García-Cardeña. Vascular endothelium, hemodynamics, and the pathobiology of atherosclerosis. Cardiovascular Pathology: The Official Journal of the Society for Cardiovascular Pathology, 22(1):9–15, February 2013. ISSN 1879-1336. doi: 10.1016/j.carpath.2012.06.006.

[9] Göran K. Hansson. Inflammation, atherosclerosis, and coronary artery disease. New England Journal of Medicine, 352(16):1685–1695, April 2005. ISSN 0028-4793, 1533-4406. doi: 10.1056/NEJMra043430. URLhttp://www.nejm.org/doi/abs/10.1056/NEJMra043430.

[10] Jiyuan Tu, Kiao Inthavong, and Kelvin Kian Loong Wong. Computational Hemodynamics - Theory, Modeling and Applications.Biological and Medical Physics, Biomedical Engineering. SpringerNetherlands : Imprint: Springer, Dordrecht, 1st ed. 2015 edition, 2015. ISBN 9789401795944.

[11] Keiichi Itatani, Shohei Miyazaki, Tokoki Furusawa, et al. New imaging tools in cardiovascular medicine: computational fluid dynamics and 4D flow MRI. General Thoracic and Cardiovascular Surgery, 65(11):611–621, November 2017. ISSN 1863-6713. doi: 10.1007/s11748-017-0834-5.

[12] Celine Souilhol, Jovana Serbanovic-Canic, Maria Fragiadaki, et al. Endothelial responses to shear stress in atherosclerosis: a novel role for developmental genes.Nature Reviews Cardiology, 17(1):52–63, January 2020. ISSN 1759-5002, 1759-5010. doi: 10.1038/s41569-019-0239-5. URLhttp://www.nature.com/articles/s41569-019-0239-5.

[13] Veronique Peiffer, Spencer J. Sherwin, and Peter D. Weinberg. Does low and oscillatory wall shear stress correlate spatially with early atherosclerosis? A systematic review. Cardiovascular Research,99(2):242–250, July 2013. ISSN 1755-3245, 0008-6363. doi:10.1093/cvr/cvt044. URLhttps://academic.oup.com/cardiovascres/article-lookup/doi/10.1093/cvr/cvt044.

[14] John Charonko, Satyaprakash Karri, Jaime Schmieg, Santosh Prabhu,and Pavlos Vlachos. In vitro, time resolved piv comparison of the effect of stent design on wall shear stress.Annals of BiomedicalEngineering, 37(7):1310–1321, July 2009. ISSN 0090-6964, 1573-9686. doi: 10.1007/s10439-009-9697-y. URLhttp://link.springer.com/10.1007/s10439-009-9697-y.

[15] Taylor Suess, Joseph Anderson, Laura Danielson, et al. Examination of near wall hemodynamic parameters in the renal bridging stent of various stent graft configurations for repairing visceral branched aortic aneurysms. Journal of Vascular Surgery, 64(3):788–796, September 2016. ISSN 1097-6809. doi: 10.1016/j.jvs.2015.04.421.

[16] Huseyin Enes Salman, Burcu Ramazanli, Mehmet Metin Yavuz,and Huseyin Cagatay Yalcin. Biomechanical investigation of disturbed hemodynamics-induced tissue degeneration in abdominal aortic aneurysms using computational and experimental techniques.Frontiers in Bioengineering and Biotechnology, 7:111, 2019. ISSN2296-4185. doi: 10.3389/fbioe.2019.00111.

[17] Jinah Hwang, Aniket Saha, Yong Chool Boo, et al. Oscillatory Shear Stress Stimulates Endothelial Production of O2 - from p47-dependent NAD(P)H Oxidases, Leading to Monocyte Adhesion.Journal of Biological Chemistry, 278(47):47291–47298, November2003. ISSN 00219258. doi: 10.1074/jbc.M305150200. URLhttps://linkinghub.elsevier.com/retrieve/pii/S0021925820760242.

[18] George P. Sorescu, Hannah Song, Sarah L. Tressel, et al. Bone Morphogenic Protein 4 Produced in Endothelial Cells by Oscillatory Shear Stress Induces Monocyte Adhesion by Stimulating Reactive OxygenSpecies Production From a Nox1-Based NADPH Oxidase. Circulation Research, 95(8):773–779, October 2004. ISSN 0009-7330, 1524-4571. doi: 10.1161/01.RES.0000145728.22878.45. URLhttps://www.ahajournals.org/doi/10.1161/01.RES.0000145728.22878.45.

[19] S. Glagov, C. Zarins, D. P. Giddens, and D. N. Ku. Hemodynamics and atherosclerosis. insights and perspectives gained from studies of human arteries.Archives of Pathology & Laboratory Medicine, 112(10):1018–1031, October 1988. ISSN 0003-9985.

[20] Amanda Sampaio Storch, Helena Naly Miguens Rocha, Vinicius Pacheco Garcia, et al. Oscillatory shear stress induces hemostatic imbalance in healthy men.Thrombosis Research, 170:119–125, October 2018. ISSN 00493848. doi: 10.1016/j.thromres.2018.08.019. URLhttps://linkinghub.elsevier.com/retrieve/pii/S0049384818304791.

[21] Peter F. Davies. Hemodynamic shear stress and the endothelium in cardiovascular pathophysiology.Nature Clinical Practice. Cardiovascular Medicine, 6(1):16–26, January 2009. ISSN 1743-4300. doi:10.1038/ncpcardio1397.

[22] Ayla Hoogendoorn, Annette M. Kok, Eline M. J. Hartman, et al.Multidirectional wall shear stress promotes advanced coronary plaque development: comparing five shear stress metrics.CardiovascularResearch, 116(6):1136–1146, May 2020. ISSN 1755-3245. doi:10.1093/cvr/cvz212.

[23] Monika Colombo, Yong He, Anna Corti, et al. Baseline local hemodynamics as predictor of lumen remodeling at 1-year follow-up in stented superficial femoral arteries. Scientific Reports, 11(1):1613, January 2021. ISSN 2045-2322. doi: 10.1038/s41598-020-80681-8.

[24] D. N. Ku, D. P. Giddens, C. K. Zarins, and S. Glagov. Pulsatile Flow and atherosclerosis in the human carotid bifurcation. Positive correlation between plaque location and low oscillating shear stress. Arteriosclerosis (Dallas, Tex.), 5(3):293–302, June 1985. ISSN 0276-5047. doi: 10.1161/01.atv.5.3.293.

[25] Yiemeng Hoi, Bruce A Wasserman, Yuanyuan J Xie, et al. Characterization of volumetric flow rate waveforms at the carotid bifurcations of older adults. Physiological Measurement, 31(3):291–302, March 2010. ISSN 0967-3334, 1361-6579.doi: 10.1088/0967-3334/31/3/002. URLhttps://iopscience.iop.org/article/10.1088/0967-3334/31/3/002.

[26] Claudio Chiastra, Stefano Morlacchi, Diego Gallo, Umberto Morbiducci, Rubén Cárdenes, Ignacio Larrabide, and Francesco Migliavacca. Computational fluid dynamic simulations of image-based stented coronary bifurcation models.Journal of the Royal Society,Interface, 10(84):20130193, July 2013. ISSN 1742-5662. doi:10.1098/rsif.2013.0193.

[27] Ronny Amaya, Limary M. Cancel, and John M. Tarbell. Interaction between the stress phase angle (spa) and the oscillatory shear index(osi) affects endothelial cell gene expression.PloS One, 11(11):e0166569, 2016. ISSN 1932-6203. doi: 10.1371/journal.pone.0166569.

[28] John J. Asiruwa, Aaron M. Propst, and Stephen P. Gent. AssessingNear-Wall Hemodynamics of Blood Flow in the Left AnteriorDescending Segment of the Left Coronary Artery Using Computational Fluid Dynamics. InVolume 3: Biomedical and Biotechnology Engineering, page V003T04A023, Tampa,Florida, USA, November 2017. American Society of MechanicalEngineers. ISBN 9780791858363. doi: 10.1115/IMECE2017-71432. URLhttps://asmedigitalcollection.asme.org/IMECE/proceedings/IMECE2017/58363/Tampa,%20Florida,%20USA/263594.

[29] Yiannis S. Chatzizisis, Ahmet Umit Coskun, Michael Jonas, Elazer R. Edelman, Charles L. Feldman, and Peter H. Stone. Role of endothelial shear stress in the natural history of coronary atherosclerosis and vascular remodeling: molecular, cellular, and vascular behavior. Journal of the American College of Cardiology, 49(25):2379–2393, June2007. ISSN 1558-3597. doi: 10.1016/j.jacc.2007.02.059.

